# High-SNR Whole-Brain Mesoscale Diffusion MRI Using Rotating-View Acquisitions (ROVER-dMRI)

**DOI:** 10.64898/2026.09.25.754394

**Authors:** Qiang Liu, Yunqi Wang, Yiang Pan, Xingwang Yong, Yuting Chen, Hwihun Jeong, Nan Wang, Erpeng Dai, Jarrett Rushmore, Carl-Fredrik Westin, Lauren J. O’Donnell, Berkin Bilgic, Kawin Setsompop, Deniz Erdogmus, Lipeng Ning, Yogesh Rathi

## Abstract

**Purpose:** We aim to acquire and reconstruct high signal-to-noise ratio (SNR), whole-brain, mesoscale (isotropic submillimeter) diffusion MRI (dMRI) in vivo in a clinically feasible scan time.

**Methods:** We present ROVER-dMRI, a unified rotating-view acquisition and super-resolution reconstruction framework. Thick slices (up to 10 mm) acquired across many rotated orientations improve the base SNR, and a shared multi-resolution hash-encoded implicit neural representation (INR) with per-direction MLP head and a physics-informed, Bloch-simulated slice-profile forward model reconstructs these views into a continuous isotropic volume (up to 13× super-resolution) in a self-supervised manner. ROVER-dMRI was evaluated in vivo on human brain (0.78 mm isotropic and 0.5 mm isotropic) and monkey brain (0.6 mm isotropic) across MRI platforms with varying gradient performance, and benchmarked against gSlider-SMS and least-squares super-resolution reconstruction (LS-SRR).

**Results:** At 0.78 mm isotropic resolution, ROVER-dMRI diffusion-weighted images achieved higher SNR than the vendor product sequence, whose images were limited by the noise floor, while its directionally encoded color (DEC) maps showed lower noise than gSlider. When reducing the number of views from 20 to 13 (reducing the scan time by 65%), ROVER-dMRI acquisition/reconstruction preserved all fine details in the fiber orientations. In the monkey brain, its DEC map matched a 60-direction gSlider acquisition at only 40% of the scan time. ROVER-dMRI preserved coherent fiber-orientation-distribution (FOD) structure, including crossing fibers, and recovered fine details in the cerebellum at 0.5 mm isotropic, that are not visible at 0.78 mm resolution. Compared to state-of-the-art 0.5 mm methods, ROVER-dMRI reduced scan time by 73% while acquiring more diffusion directions.

**Conclusion:** ROVER-dMRI offers a practical route to high-SNR whole-brain mesoscale diffusion imaging in vivo, with the pulse sequence and reconstruction pipeline released as open source.

## 1 INTRODUCTION

Diffusion MRI (dMRI) enables noninvasive characterization of brain microstructure and connectivity. Achieving isotropic submillimeter (mesoscale) resolution reveals fine-scale neural architectures that are obscured by severe partial-volume effects at standard clinical resolutions (e.g., 2 mm) ^1^. At mesoscale spatial resolution, dMRI improves connectivity-based cortical parcellation ^2^, detects silent microinfarcts ^3^, and precisely delineates short association U-fibers and highly curved white-matter pathways ^4,5^, thereby mitigating tractography “gyral bias” at the gray–white matter boundary ^6^. It also allows mapping intricate subcortical structures crucial for deep-brain-stimulation targeting ^7^. Achieving such resolution in vivo, however, poses a fundamental physical challenge: because whole-brain submillimeter dMRI typically relies on two-dimensional echo-planar imaging (2D-EPI), the drastic reduction in voxel size inherently causes severe signal-to-noise ratio (SNR) starvation.

To overcome the inherent SNR starvation of sub-millimeter dMRI, both advanced acquisition and reconstruction strategies have been developed ^8,9^. On the acquisition side, 3D-volumetric and multi-slab sequences improve SNR efficiency ^10,11,12,13^ but remain susceptible to shot-to-shot phase variations, which require either self-navigated readouts or navigators. As a 2D sequence, gSlider ^14,15^ uses thick-slab RF encoding to boost the SNR at the cost of slab-boundary artifacts, which further require 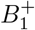 and *T*_1_ correction. Building on gSlider-SMS ^14,16^ and blip up-down acquisition (BUDA) ^16^, PRIME (phase-reversed interleaved multi-echo acquisition) was proposed ^17^, and on the high-performance head-only Connectome 2.0 scanner, high-fidelity mesoscale DWI at 0.49 mm isotropic resolution was obtained in vivo. To address the limitations of acquisition-centric methods, computational super-resolution reconstruction (SRR) have been increasingly utilized. SRR reconstructs a single high-isotropic-resolution volume from multiple low-resolution, high-SNR volumes, typically acquired with sub-voxel spatial shifts or rotated slice geometries, to exploit the complementary spatial information they sample ^8,18,9^. This recovery is traditionally formulated as an inverse problem and solved via regularized least-squares (LS-SRR). A recent prominent example is Romer-EPTI ^19^, which combines rotating-view thick-slice acquisitions with Echo-Planar Time-resolved Imaging to achieve distortion-free, motion-robust SRR. However, diffusion MRI acquisition using Romer-EPTI involves substantially long acquisition times: for instance, at 0.5 mm isotropic resolution, capturing 25 diffusion directions on a Prisma scanner takes approximately 78 min.

While such rotating-view SRR approaches generally work well when the Nyquist sampling criterion is satisfied, the underlying standard LS-SRR solver fundamentally operates on fixed, discrete voxel grid and relies on predefined mathematical priors, which can be inadequate in high noise scenarios or when undersampling is desired to reduce scan the long scan times.

To overcome these limitations, implicit neural representations (INRs) ^20,21^ have emerged as a powerful alternative. Rather than storing an image on a discrete grid, an INR maps spatial coordinates directly to signal intensities, acting as a continuous, flexible prior that learns the anatomy directly from the sampled data. Because this learning is self-supervised, it requires no externally labeled training datasets. Traditionally, INRs model this coordinate-to-signal mapping with a single global multi-layer perceptron (MLP), but this design scales poorly to high-resolution isotropic volumes: such networks struggle to capture high-frequency local details and require prohibitively long reconstruction times, since the very large MLPs needed for global parameterization suffer from diminishing representational capacity. Multi-resolution hash encoding ^22,23^ addresses this bottleneck by embedding spatial coordinates into compact, hierarchical hash tables before passing them to a much smaller MLP, capturing both global context and fine anatomical detail with far greater efficiency.

Building on this computational advance, rotating-view super-resolution MRI (ROVER-MRI) ^24^ established a powerful synergy between multi-view acquisition and continuous reconstruction. On the acquisition side, ROVER-MRI employs thick 2D slices across rotated orientations to maximize SNR. While thick slices inherently lack high-frequency detail in the through-plane direction, they retain high spatial resolution in-plane. By rotating the acquisition angles, the high-resolution in-plane data of one view naturally intersects and compensates for the low-resolution through-plane dimension of another, capturing strictly complementary spatial information. During reconstruction, it explicitly leverages multi-scale hash encoding to efficiently map the complex, intersecting spatial coordinates of these rotated slices, allowing the INR to seamlessly aggregate diverse geometric projections into a unified, continuous isotropic volume. This approach has achieved 5 to 11× resolution enhancement in high-SNR structural imaging (e.g., *T*_2_-SPACE ^25^). To date, however, this framework has been demonstrated only on high-SNR structural data; whether such INR-based reconstruction can be extended to mesoscale diffusion MRI, where the SNR is significantly lower and the data encompass complex angular dimensions, remains unexamined.

Building on the success of ROVER-MRI in structural imaging, in this work, we extend the continuous representation capabilities of INRs to the dMRI domain, addressing the ultra-low-SNR challenge that has long constrained in vivo mesoscale dMRI. Whereas prior efforts to reach submillimeter diffusion resolution have relied on complex acquisition sequences, specialized encoding schemes, or high-performance gradient systems, we propose ROVER-dMRI (rotating-view super-resolution diffusion MRI), a unified acquisition and continuous reconstruction framework for ultra-high-resolution diffusion imaging. Our approach leverages thick slices (up to 10 mm) to dramatically elevate the base SNR, then combines the complementary information across these rotated, SNR-rich acquisitions using a multi-scale hash-encoded INR to recover thin-slice isotropic volumes with up to a 13× super-resolution factor. Specific to dMRI, which acquires many diffusion directions, we reconstruct several gradient-encoded volumes jointly with a multi-head INR that shares a single hash encoding across directions and adds a lightweight per-direction output MLP head, pooling their common anatomy to suppress noise while preserving per-direction contrast. The main contributions of this work are as follows:

- **Continuous mesoscale framework with physics-informed slice profile correction**. We develop ROVER-dMRI, a multi-scale implicit neural representation framework tailored specifically for diffusion data. Moving beyond the simplified formulation of the original ROVER-MRI that ignored slice profiles, our framework incorporates a physics-informed forward model that accurately reflects actual signal formation. Specifically, we aggregate the continuous sub-slice predictions using normalized slice-profile weights derived directly from Bloch simulations of the RadioFrequency (RF) pulses. This ensures accurate mapping of the contribution of the signal from each thin-slice to the thick-slice acquisition. Additionally, we enhance the INR architecture to allow artifact-free reconstruction using much fewer views than required by Nyquist criteria, thereby reducing the scan time. We test ROVER-dMRI for in vivo mesoscale dMRI across multiple MRI platforms with differing gradient performance and at various target resolutions, down to 0.5 mm isotropic.
- **Superior microstructural fidelity and robust FOD estimation**. We systematically validate the microstructural fidelity of ROVER-dMRI against gSlider-SMS acquisition. Using diffusion-tensor-derived directionally encoded color (DEC) maps, we demonstrate the ability of our framework to preserve detailed white-matter anatomy, in highly complex regions such as the cerebellum. Furthermore, we evaluate fiber orientation distribution (FOD) maps to assess the accurate recovery of complex crossing-fiber structures. Crucially, we analyze the robustness of our framework to data undersampling by comparing reconstructions from a 20-view acquisition (which meets the Nyquist sampling rate) against a reduced 13-view subset, illustrating the impact of view reduction and the capability of ROVER-dMRI to maintain high microstructural fidelity even with accelerated scan times.
- **Automated open-source diffusion acquisition sequence**. While ROVER-dMRI acquisition can be setup manually using product sequences (with a separate sequence setup for each rotating view), we designed and implemented a custom multi-view diffusion pulse sequence using the open-source Pulseq framework ^26^ as well. This enables the automated, continuous acquisition of multiple rotated views within a single scan protocol, eliminating the operational friction of manual slice repositioning. To foster transparency and collabo nrative sequence development, the source code for both the multi-view diffusion pulse sequence and the INR reconstruction pipeline is made publicly available at https://github.com/QiangLiu0310/ROVER-dMRI.git. Example datasets of this study are available from the corresponding author upon request.

## METHODS

### 2.1 ROVER-dMRI framework

Figure 1 gives an overview of the ROVER-dMRI framework. We reconstruct super-resolution diffusion-weighted images from multiple rotated thick-slice acquisitions using a coordinate-based implicit neural representation (INR). The *D* diffusion-weighted volumes (gradient directions) are reconstructed jointly by a single network (Section 2.1.3). The pipeline has three stages: (1) coordinate mapping, which registers every view into a common world frame; (2) a novel multi-resolution multi-hash encoding followed by a multi-head MLP, in which the hash features are shared across all diffusion directions and one lightweight head per direction regresses the thin-slice signal at *M* sub-slice positions per thick slice; and (3) a physics-informed loss that aggregates those sub-slice predictions through the measured slice profile and is summed over all directions before comparing to the acquired thick-slice data.

**FIGURE 1.**
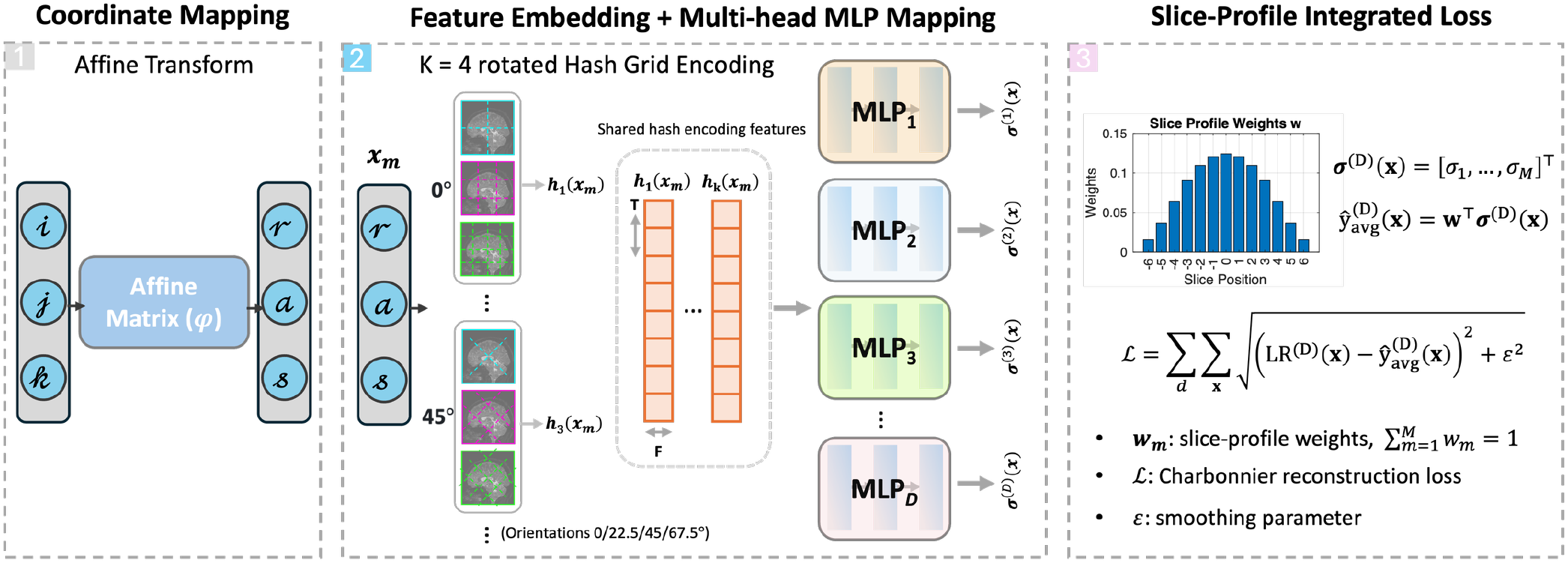
Overview of the ROVER-dMRI framework. Rotated thick-slice low-resolution volumes are mapped into a common world frame by the affine transforms ***φ***_*v*_ and embedded by a multi-resolution hash encoder ***h***(*·*) whose features are shared across all diffusion directions. A multi-head MLP with one head 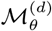 per diffusion direction *d* decodes the shared feature into the thin-slice signal ***σ***^(*d*)^(***x***) of that direction at *M* sub-slice positions. The per-direction sub-slice predictions are aggregated through the slice-profile weights ***w*** and trained against the measured data with a physics-informed loss summed over all directions.

#### 2.1.1 Coordinate mapping

The views are first placed in a common spatial reference frame. Let LR_*v*_ be the *v*-th low-resolution volume, acquired at rotation angle *θ*_*v*_ with integer voxel indices (*i, j, k*). Each index triplet is mapped to real-world Right–Anterior–Superior (RAS) coordinates (*r, a, s*) by the view-specific affine transform ***φ***_*v*_ :

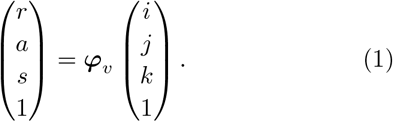

The affine ***φ***_*v*_ encodes the rotation and translation of view *v*. For a rotation by angle *θ*_*v*_ about the phase-encoding axis,

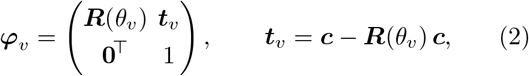

where ***R***(*θ*_*v*_) is the rotation matrix and the translational term ***t***_*v*_ recenters the rotation on the FOV center ***c*** = (*r*_*c*_, *a*_*c*_, *s*_*c*_)^*⊤*^.

Only the coordinates are transformed; the measured voxel intensities are left unchanged and serve as the supervision signal during training. Because each thick-slice voxel integrates signal across the slice-select direction, we represent it by *M* evenly spaced sub-slice sample positions along that direction. This yields, for each thick-slice voxel, a set of continuous coordinates {***x***_*m*_, *m* = 1, …, *M*}. All coordinates are normalized to [0, 1] before feature embedding.

#### 2.2.2 Feature embedding with rotated multi-resolution hash grids

Each normalized coordinate ***x***_*m*_ is encoded by a multi-resolution hash grid implemented in the Tiny CUDA Neural Networks (TCNN) framework ^27^, whose fused CUDA kernels make hashing, lookup, and interpolation fast enough to train on a single GPU.

The encoder ***h***(·) maps ***x***_*m*_ to a feature vector by querying *L* resolution levels at once. Each level *l* holds a hash table of size *T* storing *F* -dimensional features at grid vertices. Level resolutions follow a geometric progression between the coarsest and finest grids:

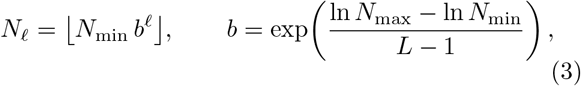

where *b* is the per-level scale factor, the constant geometric ratio by which the grid resolution grows from one level to the next. At level *l*, ***x***_*m*_ is scaled to that grid, the enclosing cell is found, and its eight corner vertices are mapped to table entries by a spatial hash:

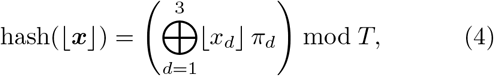

where *π*_*d*_ are large primes and ⊕ is the bitwise XOR. The eight corner features are trilinearly interpolated and the *L* levels concatenated into a per-grid feature ***h***_*k*_(***x***) = [***f*** ^(1)^(***x***), …, ***f*** ^(*L*)^(***x***)].

A single hash grid aligns its lattice with the reconstruction-grid axes. Because the rotating thick-slice acquisition constrains only a subspace of the isotropic volume, such an axis-aligned lattice tends to fill the unconstrained directions with structure along its own axes, which appears as fine line artifacts in the reconstruction; grid-based encodings are known to carry this axis-aligned bias ^28^. To suppress it, we replace the single grid with an ensemble of *K* cubic hash grids whose lattices are rotated to distinct orientations. Grid *k* encodes the coordinate rotated by a fixed matrix ***R***_*k*_ (with ***R***_1_ = ***I***), and the per-grid features are concatenated, together with the raw coordinate, into the shared encoding

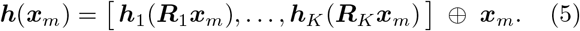

Combining the *K* differently oriented grids averages down the axis-aligned stripes while retaining the shared, data-constrained anatomy. We use *K* = 4 grids at 0*/*22.5*/*45*/*67.5^*°*^ with *F* = 1 feature per grid, and cap the finest grid resolution *N*_max_ at the data Nyquist resolution set by the acquired in-plane sampling, dropping any level whose resolution *N*_*l*_ would exceed it. Encoding parameters (*K, L, T, F, N*_min_, and the per-level scale *b*) are reported in Table 1.

**TABLE 1.** Network and reconstruction hyperparameters per dataset.

| Parameter | 0.78 mm iso | 0.6 mm iso (monkey) | 0.5 mm iso |
| --- | --- | --- | --- |
| Number of rotated grids ($K$) | 4 | 1 | 4 |
| Number of resolution levels ($L$) | 7 | 9 | 8 |
| Hash table size ($T$) | $2^{25}$ | $2^{26}$ | $2^{25}$ |
| Feature dimensions per entry ($F$) | 1 | 2 | 1 |
| Per-level scale ($b$) | 1.6 | 1.5 | 1.6 |
| Coarsest resolution ($N_{\min}$) | 16 | 16 | 16 |
| Charbonnier parameter $\varepsilon$ | 1e-1 | 1e-3 | 1e-1 |

#### 2.1.3 Multi-head MLP regression of the thin-slice signal

We evaluated ROVER-dMRI at three target isotropic resolutions: 0.78 mm (human brain), 0.6 mm (in-vivo monkey brain), and 0.5 mm (human brain).

For all three datasets, the *D* diffusion-weighted volumes are reconstructed jointly by a multi-head MLP. The shared multi-resolution hash features ***h***(***x***_*m*_) is are estimated jointly per coordinate and shared across all diffusion directions; and decoded individually by *D* lightweight heads, one per diffusion-weighted volume (gradient direction). In the actual implementation, *D* was set to 6 to reduce noise and improve the estimation of the anatomy. Each head 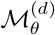 is a compact MLP (two hidden layers, each with 128 neurons, ReLU activations for the 0.78 mm and 0.6 mm datasets; three hidden layers, each with 64 neurons, ReLU activations for the 0.5 mm dataset). that maps the shared feature to a scalar intensity. Evaluating head 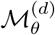 at each of the *M* sub-slice coordinates gives the thin-slice prediction of direction *d* at sub-slice *m*:

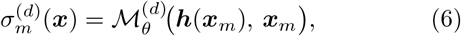

for *m* = 1, …, *M* and *d* = 1, …, *D*, where ***h***(***x***_*m*_) is the multi-resolution hash feature and ***x***_*m*_ is the auxiliary positional input. Stacking the *M* outputs of head 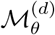 gives its sub-slice signal vector 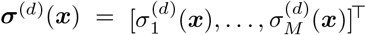 along the slice-select direction, which Section 2.1.4 aggregates through the slice profile.

The shared encoding is updated by gradients from all *D* directions, whereas each head is trained on the data of its own direction. The encoding and heads are implemented in TCNN and PyTorch and optimized jointly. At inference, the shared encoding and all *D* heads are evaluated on a uniform grid at the target isotropic spacing to produce the *D* super-resolved diffusion volumes in a single pass. Because the *b* = 0 volume differs markedly in contrast from the diffusion-weighted volumes, it is excluded from this joint multi-head reconstruction and instead reconstructed with its own separate network, trained independently on the *b* = 0 data alone.

#### 2.1.4 Physics-informed loss via slice-profile aggregation

To stay consistent with the thick-slice measurements, for each direction *d* the *M* sub-slice predictions are combined using slice-profile weights ***w*** = [*w*_1_, …, *w*_*M*_]^*⊤*^ obtained from a Bloch simulation of the RF excitation profile, discretized at the *M* sub-slice positions and normalized so that 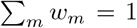. The modeled thick-slice signal of direction *d* at ***x*** is the profile-weighted sum

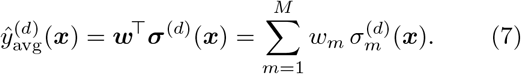

Unlike the equal-weight aggregation (*w*_*m*_ = 1*/M*) of the original ROVER-MRI, this weighting follows the simulated excitation profile, with higher weight near the slice center and lower toward the edges, and so better matches the signal formation of the sequence; the weights ***w*** are shown in Figure 1.

The training loss is a data-consistency term, the Charbonnier loss, summed over all directions and sampled coordinates:

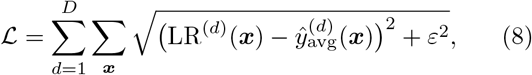

where LR^(*d*)^ is the measured low-resolution data of direction *d*. The Charbonnier loss is a differentiable, outlier-robust approximation to the *L*_1_ norm with smoothing parameter *ε*. No explicit total-variation or other spatial regularization term is used.

### 2.2 Data acquisition

As mentioned in Section 2.1.3, we evaluated ROVER-dMRI at three target isotropic resolutions, comparing against gSlider acquisitions under matched conditions where available. All studies were IRB-approved with written informed consent from two human participants (one male and one female). Acquisition parameters are listed in Table 2. The rotation axis was aligned with the phase-encoding direction for every dataset: anterior– posterior (0.78 mm), head–foot (0.6 mm monkey), and anterior–posterior (0.5 mm), to keep geometric distortion consistent across views.

**TABLE 2.** Acquisition parameters for all datasets. ROVER-dMRI is compared against gSlider under matched conditions at 0.78 mm (human, Siemens Prisma) and 0.6 mm (monkey, Siemens Connectome 2.0); the 0.5 mm protocol (Siemens Cima) used a custom Pulseq sequence.

| Parameter | 0.78 mm iso |  | 0.6 mm iso (monkey) |  | 0.5 mm iso |
| --- | --- | --- | --- | --- | --- |
|  | ROVER-dMRI | gSlider-SMS | ROVER-dMRI | gSlider | ROVER-dMRI |
| Scanner & field | Siemens Prisma 3T |  | Siemens Connectome 2.0 3T |  | Siemens Cima 3T |
| Head coil (channels) | 32 |  | 16 |  | 32 |
| Sequence type | Product | C2P | Product | C2P | Pulseseq |
| FOV (mm<sup>3</sup>) | 200 × 200 × 200 | 220 × 220 × 129 | 200 × 132 × 108 | 200 × 138 × 57 | 200 × 200 × 168 |
| Matrix (RO×PE) | 256 × 256 | 290 × 290 | 332 × 206 | 332 × 230 | 400 × 400 |
| Spatial res. (mm<sup>3</sup>) | 0.78<sup>3</sup> | 0.76<sup>3</sup> | 0.6<sup>3</sup> | 0.6<sup>3</sup> | 0.5<sup>3</sup> |
| Slice thickness (mm) | 10 | 3.8 | 6 | 3 | 6 |
| Number of slices | 20 | 34 | 18 | 19 | 28 |
| TR (ms) | 2000 | 3500 | 2500 | 3000 | 2200 |
| TE (ms) | 80 | 87 | 68 | 80 | 74 |
| In-plane acceleration | 3 | 3 | 2 | 2 | 2 (3-shot) |
| Partial Fourier | 6/8 | 6/8 | 6/8 | 6/8 | 6/8 |
| Multiband factor | 2 | 2 | 1 | 1 | 2 |
| Rotation views, angle | 20, 9° | 5 (RFs) | 15, 12° | 5 (RFs) | 12, 15° |
| $b$-values (s/mm<sup>2</sup>) | 0 / 1000 | 0 / 1000 | 0 / 1000 | 0 / 1000 | 0 / 1000 |
| Diffusion directions | 60 | 60 | 30 | 60 | 30 |
| Echo spacing (ms) | 1.08 | 1.13 | 0.95 | 0.95 | 1.54 |
| Total scan time (min) | 40:40 | 17:47 | 28:00 | 68:48 | 43:34 |

Conventionally, the number of rotated views needed to constrain the isotropic reconstruction follows the geometric framework for super-resolution MRI ^29,24^: the theoretical minimum number of rotations *N*_*R*_ should satisfy

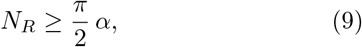

where *α* is the super-resolution factor, the ratio of the slice thickness to the in-plane (isotropic) resolution.

For the 0.78 mm and 0.6 mm datasets, rotating-view low-resolution volumes were acquired with the vendor diffusion-weighted EPI sequence, rotating the FOV manually between repeated acquisitions. To reach 0.5 mm in-plane resolution with automated rotation, we implemented a custom 3-shot diffusion-weighted EPI sequence in Pulseq ^26,30,31^, using the Pulseq rotation extension (Pulseq v1.5.1) to acquire all rotated views within a single sequence run.

### 2.3 Data processing and super-resolution reconstruction

For the 0.78 mm and 0.6 mm datasets, vendor-reconstructed DICOMs (ROVER-dMRI and gSlider) were exported and converted to NIfTI; distortion was then corrected with FSL TOPUP, followed by eddy-current and motion correction with FSL Eddy ^32^. For the 0.5 mm dataset, raw *k*-space was converted to the ISMRM-RD format ^33^ and reconstructed offline with multi-shot DPG for Nyquist ghost correction ^34,35,36^and S-LORAKS ^37^; the result was converted to NIfTI.

Super-resolution reconstruction used the ROVER-dMRI framework of Section 2.1. Intensities were normalized to [0, 1] before training. The network was optimized with Adam (learning rate 10^*−*4^) using the Charbonnier loss of Section 2.1.4; with the parameter *ε* set as in Table 1. Each step used an effective batch of 1 000 000 coordinates, formed from 40 000 sampled coordinates with gradient accumulation over 25 micro-batches. For comparison, conventional LS-SRR ^9^ was applied to the 0.78 mm dataset from the same low-resolution inputs.

### 2.4 Image evaluation

#### 2.4.1 Effective-resolution analysis

In EPI-based diffusion imaging, the nominal in-plane resolution overstates the true resolving capability due to a blur that is intrinsic to the in-plane EPI read-out itself, arising before the INR-based super-resolution reconstruction comes into play: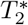 decay during the readout attenuates high spatial frequencies in *k*-space and broadens the point-spread function (PSF), predominantly along the phase-encode (PE) direction. Following Feizollah and Tardif ^38^, we estimated the effective resolution of the 0.5 mm protocol from the 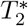 -weighted PE PSF. We applied the modulation transfer function 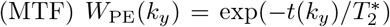, where *t*(*k*_*y*_) is the acquisition time of each PE line derived from the 3-shot echo train (≈ 76 ms per-shot readout), and took the effective resolution as the FWHM of the normalized-magnitude PSF obtained by inverse Fourier transform on a zero-padded grid. Partial-Fourier lines were restored by conjugate (Hermitian) symmetry to match the in-vivo S-LORAKS reconstruction. The fully sampled readout direction, which carries negligible 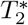 apodization over its short (∼1.54 ms) readout, was modeled as the ideal band-limited (nominal-resolution) sinc; the 2D PSF was formed as the outer product of the readout and PE responses. The analysis was run for gray matter (GM) and white matter (WM) using 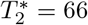 and 46 ms at 3 T, respectively ^39,40^, and repeated over nominal resolutions of 0.5 mm to 1.0 mm; see Results section (Figure S3) for more details.

#### 2.4.2 Image sharpness measurement with Laplacian variance

Quantifying the effective resolution in 3D for the proposed ROVER-dMRI method is not possible due to the non-linear nature of the reconstruction process. However, a proxy for estimating the effective resolution in 3D is to quantify the sharpness of the reconstructed DWIs. Image sharpness for the 0.78 mm isotropic dataset was quantified with the 3D Laplacian variance, a focus metric in which higher values indicate a sharper image ^41^. The standard 2D Laplacian kernel,

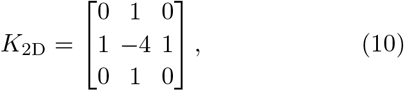

was extended to 3D as the 6-neighbor kernel spanning slices *z* − 1, *z, z* + 1,

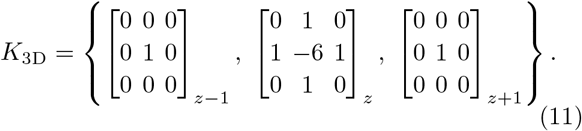

For each dataset (LS-SRR, ROVER-dMRI, and gSlider), the 3D Laplacian variance was computed over a min– max normalized 50×70×50 voxel volume centered on the cerebellum. Sharper images would imply reconstruction quality closer to the desired nominal resolution.

#### 2.4.3 Diffusion-metric comparison

Diffusion tensors were fit for all datasets, and directionally encoded color (DEC) maps were generated to assess white-matter anatomy and fiber orientation. For the 0.78 mm dataset, we additionally estimated the fiber orientation distributions (FODs) from gSlider-SMS and ROVER-dMRI using both the full 20-view acquisition and a reduced 13-view subset, to examine the effect of view reduction on reconstructed orientations and crossing-fiber structure.

## RESULTS

### 3.1 Physics-informed slice-profile aggregation

Figure 2 illustrates the effect of incorporating slice-profile aggregation into the forward model. Figure 2 (A) shows the Bloch-simulated slice profile of a 10 mm slice excited by a SINC pulse with a time–bandwidth product (TBWP) of 4. Figure 2 (B) shows the corresponding discretized and normalized sub-slice weights derived from this profile, which replace the equal weights used in the original ROVER-MRI. Figure 2 (C) compares dMRI reconstructions obtained with equal weights and with slice-profile weights, shown for both *b* = 0 and *b* = 1000 s*/*mm^2^. With equal weights, line artifacts appear throughout the brain (yellow arrows and boxes). Once slice-profile aggregation is introduced, these artifacts are removed, indicating that the physics-informed weighting better matches the actual signal formation and improves image fidelity.

**FIGURE 2.**
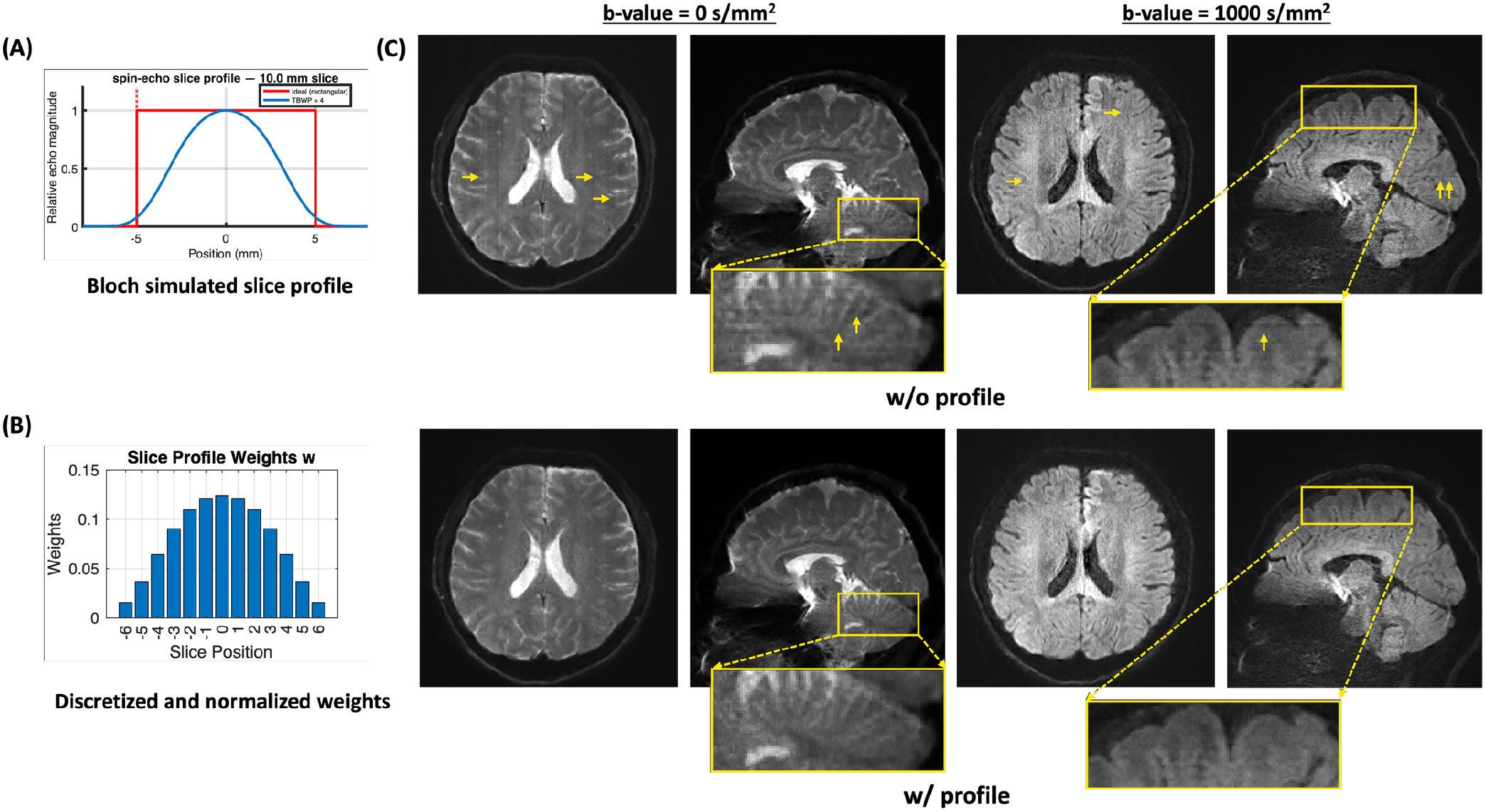
Effect of physics-informed slice-profile aggregation on dMRI reconstruction. (A) Bloch-simulated slice profile of a 10 mm slice excited by a SINC pulse with a time–bandwidth product (TBWP) of 4. (B) Discretized and normalized sub-slice weights derived from the profile in (A), used in the weighted aggregation of Equation (7); these replace the equal weights of the original ROVER-MRI by assigning higher weights near the slice center and lower weights toward the edges. (C) Comparison of dMRI reconstructions using equal weights versus slice-profile weights, shown for both *b* = 0 and *b* = 1000 s*/*mm^2^. Equal-weight aggregation produces line artifacts distributed across the whole brain (yellow arrows and boxes), which are removed once slice-profile aggregation is applied.

### 3.2 Human brain at 0.78 mm isotropic

Figure 3 compares the isotropic 0.78 mm diffusion images (*b* = 0, 1000 s*/*mm^2^) from the product sequence, gSlider-MB2, LS-SRR, and ROVER-dMRI. The scan time per volume is indicated on the left (17.5 s for gSlider-MB2 and 40 s for LS-SRR and ROVER-dMRI). Results from a faster 13-view ROVER-dMRI (26 s) are also shown, together with the corresponding voxel-wise absolute-error maps relative to the 20-view reconstruction (absolute error scaled by 5× for visualization). ROVER-dMRI achieved substantially higher SNR than the product sequence due to its intrinsically larger voxel volume, along with clearer anatomical delineation, whereas the images of the product sequence were further limited by their inherent noise floor. In contrast, gSlider-MB2 exhibited slab-boundary artifacts and elevated noise. The 13-view ROVER-dMRI preserved image detail and structure comparable to the 20-view reconstruction while nearly halving the scan time, with only minor residual differences confined to the high-noise region (i.e., the deep-brain region, as quantified by the absolute-error maps).

**FIGURE 3.**
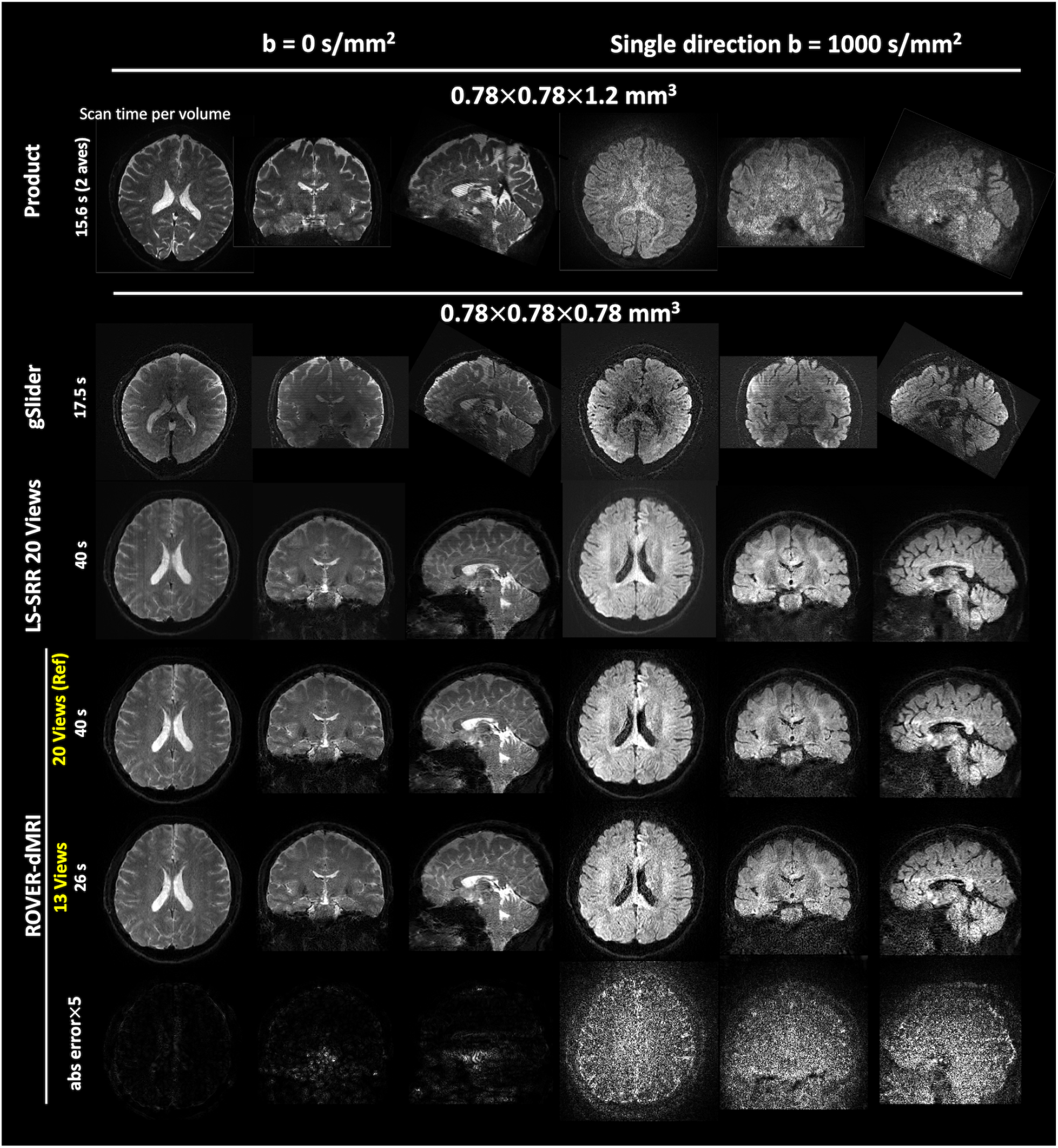
Comparison of mesoscale diffusion reconstructions across acquisition and reconstruction methods. Diffusion images from the product sequence, gSlider-MB2, LS-SRR, and ROVER-dMRI. The product sequence was acquired at 0.78 *×* 0.78 *×* 1.2 mm^3^, while gSlider-MB2, LS-SRR, and ROVER-dMRI achieved 0.78 mm isotropic resolution. (Please refer to Table 2 for the detailed scan protocol.) Representative *b* = 0 and *b* = 1000 s*/*mm^2^ images are shown, with scan times per volume indicated (17.5 s for gSlider-MB2 and 40 s for ROVER-dMRI). The rightmost columns show a faster 13-view ROVER-dMRI (26 s) and its voxel-wise absolute-error map relative to the 20-view ROVER-dMRI reconstructiosn.

Figure 4 quantifies image sharpness in the cerebellum using the 3D Laplacian variance, a standard focus metric in which a higher value indicates a sharper, less blurred image, computed over a 50 × 70 × 50 voxel volume for LS-SRR, ROVER-dMRI, and gSlider at the 0.78 mm isotropic resolution. ROVER-dMRI with 20 views achieved the highest Laplacian variance among all measurements, i.e., the sharpest images with one-voxel wide, with white matter pathways clearly visible. This analysis indicates that ROVER-dMRI preserves a sharper effective point-spread function than the other methods evaluated.

**FIGURE 4.**
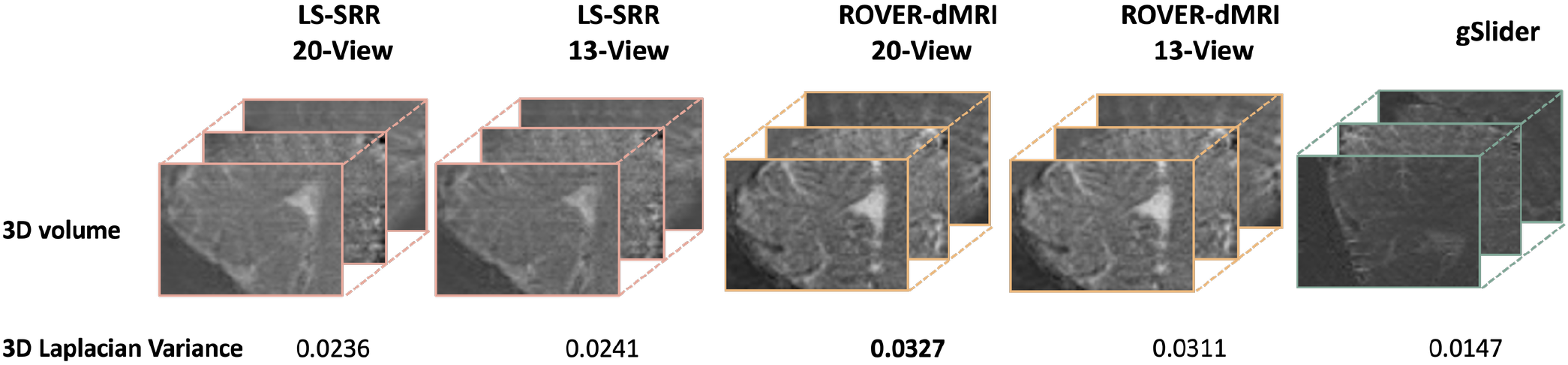
3D Laplacian variance of a 50 *×* 70 *×* 50 voxel volume in the cerebellum, comparing image sharpness across LS-SRR (20-view and 13-view), ROVER-dMRI (20-view and 13-view), and gSlider at 0.78 mm^3^ isotropic resolution. ROVER-dMRI with 20 views achieved the highest Laplacian variance (0.0327) among all measurements, compared with 0.0311 for the 13-view acquisition, 0.0236 and 0.0241 for LS-SRR with 20 and 13 views, and 0.0147 for gSlider, indicating that ROVER-dMRI preserves a sharper effective point-spread function than the other methods.

Figure S1 (Supporting Information) shows additional representative images reconstructed by LS-SRR (20-view and 13-view) and by ROVER-dMRI for the 0.78 mm dataset. The LS-SRR reconstructions exhibit line artifacts, as indicated by the arrows, and these artifacts become more severe as the number of views is reduced, whereas ROVER-dMRI images show no such line artifacts. This is consistent with the previous findings of Lyu et al. ^24^.

Figure 5 shows color-encoded FA maps (first three columns) and FODs (last column, the zoomed-in region) from gSlider and ROVER-dMRI for the 0.78 mm dataset. The gSlider DEC map is noisy, whereas, consistent with a previous study ^42^, the corona radiata is clearly resolved on the sagittal DEC map of ROVER-dMRI. In the FODs, ROVER-dMRI produces clean, anatomically consistent orientations: coherent left–right fibers in the corpus callosum (CC), superior–inferior fibers in the corticospinal tract (CST), and anterior–posterior fibers in the superior longitudinal fasciculus (SLF). These orientations are preserved when the number of views is reduced nearly by half.

**FIGURE 5.**
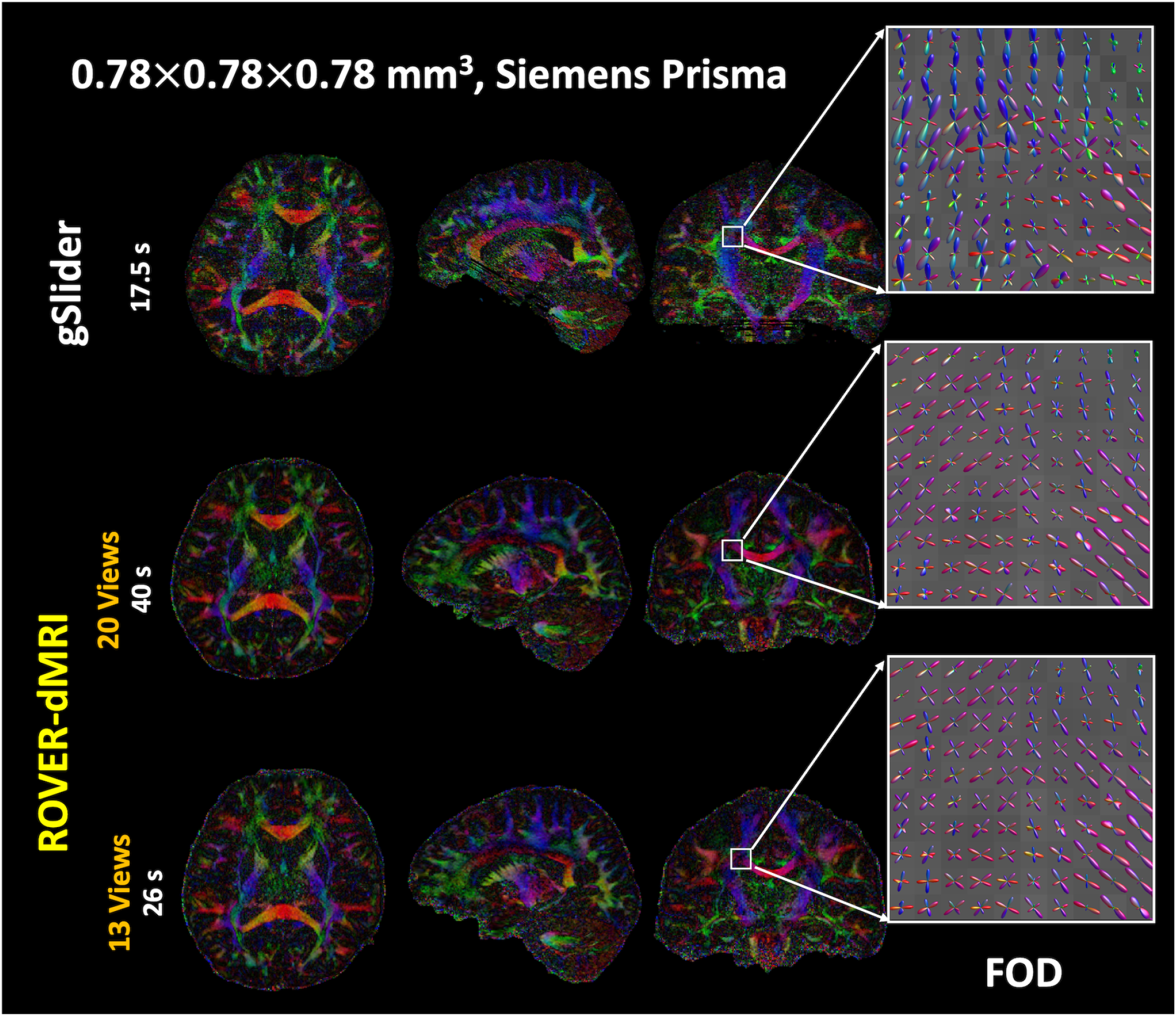
Representative color FA maps reconstructed from 60-direction DWIs using gSlider, 20-view ROVER-dMRI, and 13-view ROVER-dMRI. Corresponding FODs are shown for the corpus callosum (CC), corticospinal tract (CST), and superior longitudinal fasciculus (SLF). ROVER-dMRI yields clean and anatomically consistent orientations, with coherent left–right (red) fibers in CC, superior–inferior (blue) fibers in CST, and anterior–posterior (green) fibers in SLF. The scan time per volume is shown on the left.

Figure S2 shows DEC maps from ROVER-dMRI for the 0.78 mm dataset across varying numbers of diffusion directions and views. Using the 60-direction, 20-view reconstruction as the reference, reducing the number of diffusion directions to 30 preserves clear white-matter structure, as shown in the corona radiata and the cerebellum (zoomed insets), while reducing the number of views from 20 to 13 produces only a slight increase in noise. These results indicate that ROVER-dMRI is robust to the increased noise associated with a reduced number of diffusion directions, and that the joint reconstruction remains accurate even when the number of views (13) falls below the Nyquist sampling rate required by the reference 20-view acquisition.

### 3.3 In-vivo monkey brain at 0.6 mm isotropic

Figure 6 presents isotropic 0.6 mm dMRI data of the monkey brain. The first column shows the *T*_2_-weighted image (0.5 mm isotropic) as an anatomical reference, followed by the *b* = 0 images for the gSlider and ROVER-dMRI acquisitions. The last two columns present DEC maps computed from two acquisitions: gSlider with 60 directions (scan time 68 min 48 s, 4 averages) and ROVER-dMRI with 30 diffusion directions (28 min, a single average). On the *b* = 0 images, gSlider exhibits striping artifacts, while ROVER-dMRI yields comparable *b* = 0 image quality without these artifacts. Notably, the ROVER-dMRI DEC map closely matches that of the 60-direction gSlider acquisition while requiring only 40% of the scan time.

**FIGURE 6.**
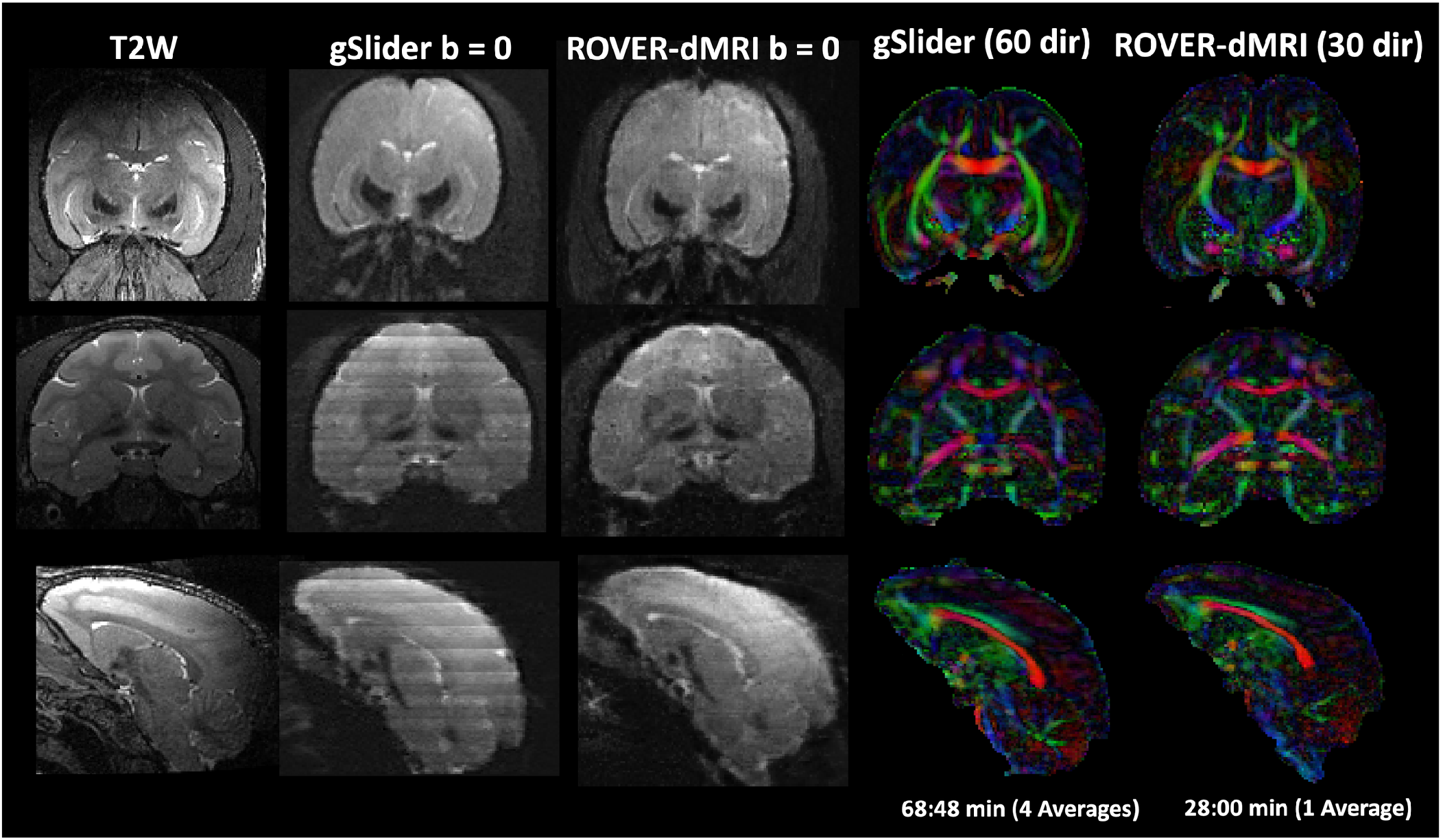
Isotropic 0.6 mm diffusion MRI of the monkey brain. The first column shows the *T*_2_-weighted anatomical reference (0.5 mm isotropic), followed by representative *b* = 0 images for the gSlider and ROVER-dMRI acquisitions. The last two columns show DEC maps computed from gSlider with 60 directions (scan time 68 min 48 s, 4 averages) and ROVER-dMRI with 30 diffusion directions (28 min, a single average). On the *b* = 0 images, gSlider shows banding artifacts that are absent in the corresponding DEC maps, whereas ROVER-dMRI provides comparable *b* = 0 image quality without such artifacts. The ROVER-dMRI DEC map closely matches that of the 60-direction gSlider acquisition while requiring only 40% of the scan time.

### 3.4 Human brain at 0.5 mm isotropic: effective resolution and mesoscale imaging

The PSF analysis (Figure S3) shows that 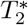 decay measurably broadens the phase-encode PSF of the 0.5 mm protocol. The 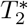 MTF is a monotonic ramp across *k*_*y*_ (Figure S3 (A)), attenuating the high-frequency edge to ∼0.3–0.5 of the *k*-space-center value (edge-to-edge ratio ≈3.2:1 for GM and ≈5.2:1 for WM). This apodization widens the PSF main lobe (Figure S3 (B)), most strongly for WM owing to its shorter 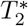. The blurring is markedly anisotropic: in the 2D PSF (Figure S3 (C)) the ringing and broadening are confined to the PE direction, whereas the rapidly sampled readout direction remains close to the ideal sinc.

At the 0.5 mm nominal resolution, the effective PE resolution was 0.66 mm for GM and 0.73 mm for WM (Figure S3 (D)), i.e. 1.3–1.5× the nominal pixel. Over the 0.5 mm to 1.0 mm range the effective resolution remained above the nominal pixel throughout and diverged from the identity line toward finer resolutions, where the longer readout increases the 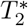 taper.

Figure 7 shows representative ROVER-dMRI images at 0.5 mm isotropic resolution in axial, coronal, and sagittal planes: a single *b* = 0 volume (top row) and a single *b* = 1000 s*/*mm^2^ diffusion-weighted volume (bottom row). Fine cortical and subcortical detail is preserved across all three orthogonal planes, with no evidence of any artifact. High SNR is preserved in the single-direction dMRI at this resolution, likely benefiting from the thick-slice acquisition and the joint reconstruction across different diffusion directions.

**FIGURE 7.**
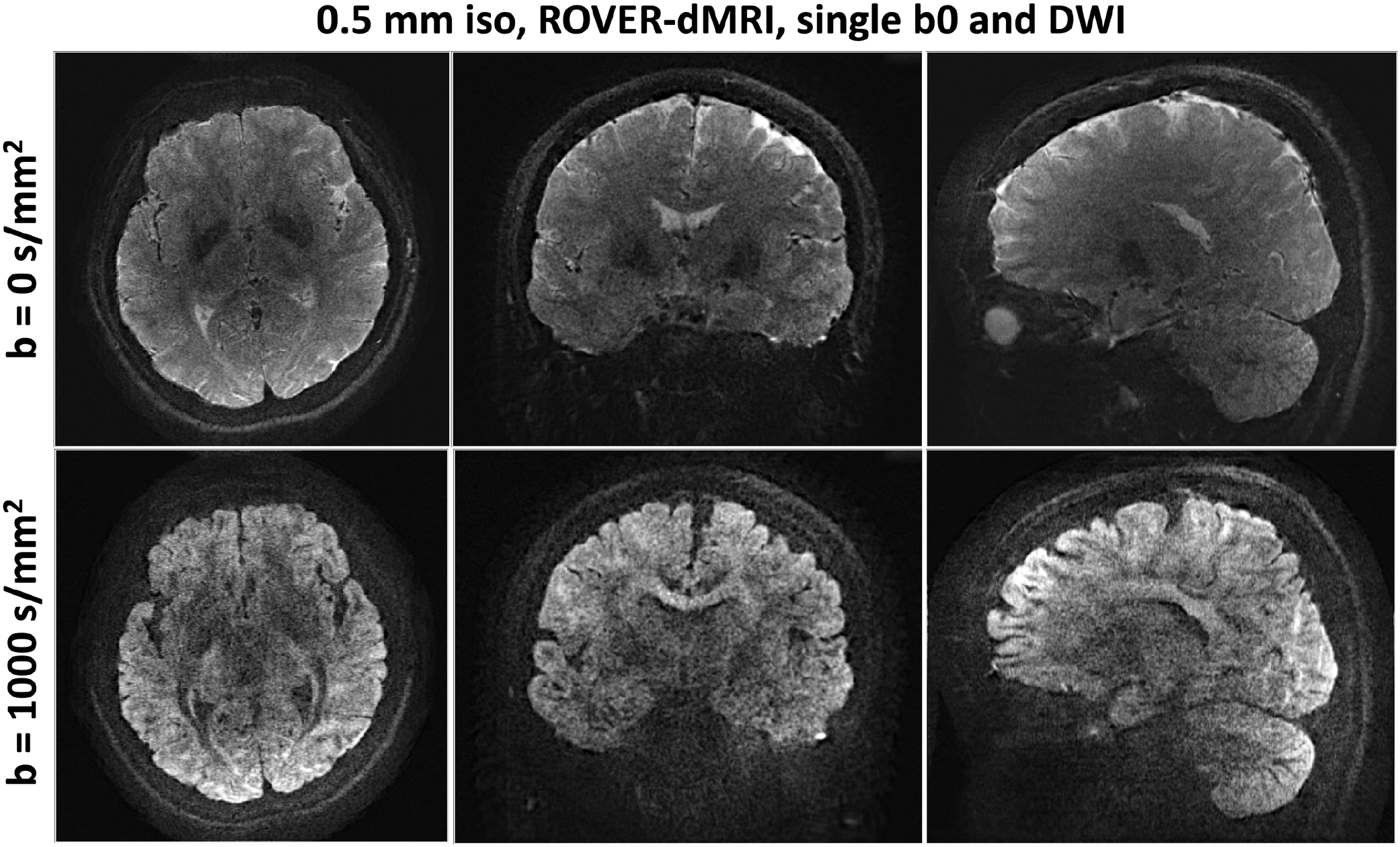
Representative ROVER-dMRI images at 0.5 mm isotropic resolution in axial, coronal, and sagittal planes. Top row: single *b* = 0 volume. Bottom row: single *b* = 1000 s*/*mm^2^ diffusion-weighted volume. Fine anatomical detail is preserved across all three planes with no evidence of significant artifact. High SNR is preserved in the single-direction dMRI at this resolution.

Figure 8 shows the 30-direction mean diffusion-weighted image (Figure 8(A)) and the corresponding DEC fractional-anisotropy map (Figure 8(B) for three representative slices in the axial view (B1, B2, and B3), and Figure 8(C) for two representative slices in the coronal view (C1 and C2)) for the same 0.5 mm isotropic ROVER-dMRI data. In the cerebellum, compared with the cFA map at 0.78 mm isotropic resolution, the 0.5 mm isotropic result reveals more anatomical detail and finer structures, as shown in the axial and coronal cFA maps and the zoomed-in regions.

**FIGURE 8.**
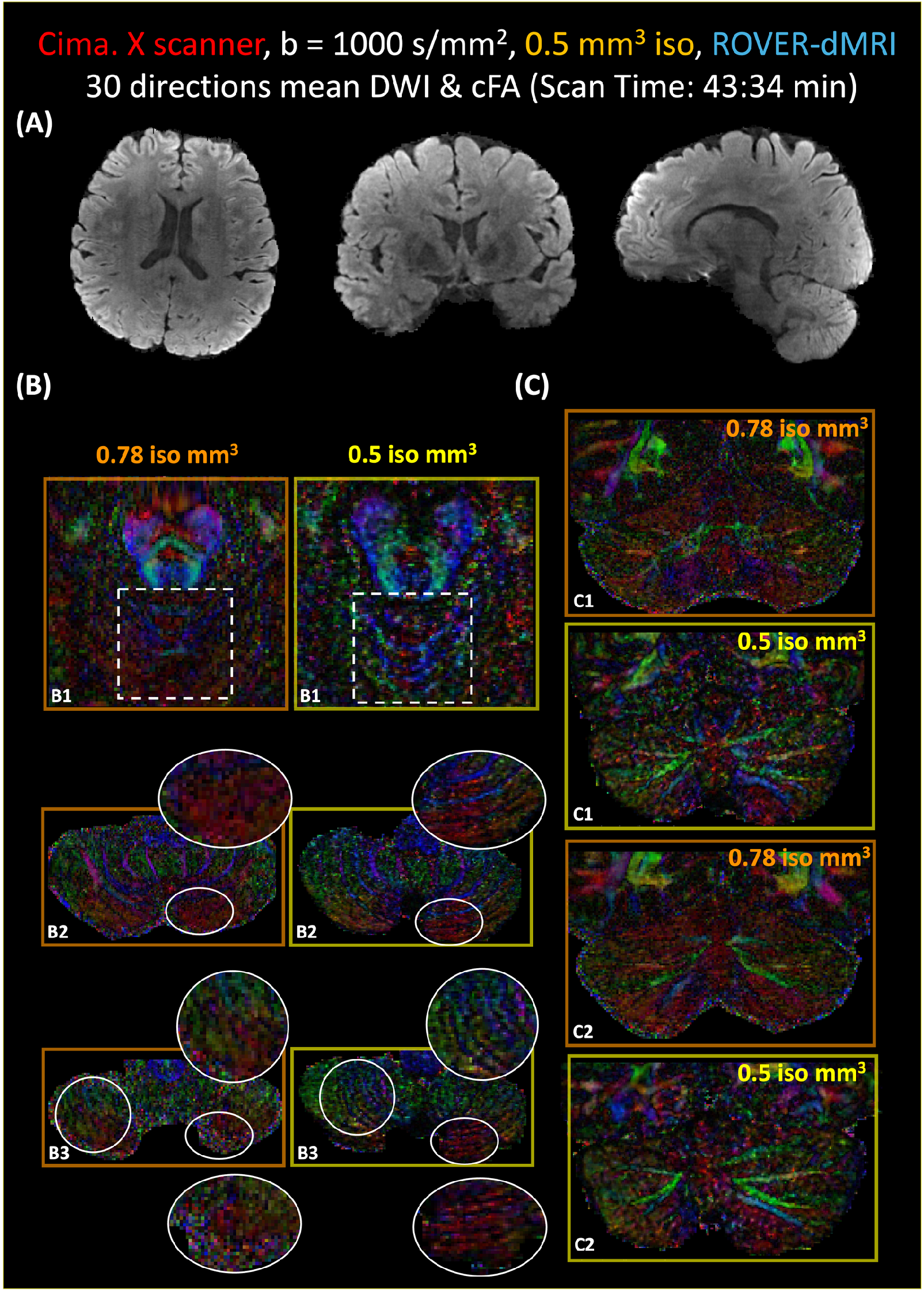
Mesoscale-averaged diffusion-weighted images (Figure 8(A)) and DEC fractional-anisotropy maps (Figure 8(B) for three representative slices (B1, B2, and B3) shown in the axial view, and Figure 8(C) for two representative slices (C1 and C2) shown in the coronal view) for *b* = 1000 s*/*mm^2^ ROVER-dMRI 0.5 mm^3^ isotropic data. High-fidelity mesoscale images were obtained from a single acquisition of 30 diffusion directions, with a total scan time of 43 min 34 s for one *b* = 0 and 30 *b* = 1000 s*/*mm^2^ volumes. The DEC fractional-anisotropy maps at 0.78 mm isotropic resolution are also shown in the cerebellum as a comparison.

## 4 DISCUSSION

In this work, we introduced ROVER-dMRI, a unified rotating-view acquisition and continuous-reconstruction framework that brings the implicit neural representation paradigm to mesoscale diffusion MRI. By acquiring thick slices across many rotated orientations, the framework elevates the base SNR that fundamentally limits in vivo submillimeter dMRI, and a multi-resolution hash-encoded INR jointly reconstructs the different diffusion gradients from these complementary views, recovering continuous isotropic volumes at up to a 13× super-resolution factor. A physics-informed forward model, aggregating the sub-slice predictions of the network through Bloch-simulated slice-profile weights rather than the equal weights of the original ROVER-MRI, removed the line artifacts associated with the simplified model and improved image fidelity. Across human (0.78 mm and 0.5 mm) and monkey (0.6 mm) experiments on scanners with differing gradient performance, ROVER-dMRI delivered higher SNR than gSlider and the vendor product sequence, preserved coherent DEC and FOD orientations including crossing-fiber populations, and retained microstructural fidelity even when the acquisition was reduced from 20 to 13 views, nearly halving the scan time.

A central advantage of ROVER-dMRI is that its SNR gain originates directly from the use of thick slices. Because the SNR of a 2D diffusion-weighted acquisition scales approximately with voxel volume, increasing the slice thickness while preserving the in-plane resolution raises the intrinsic SNR before any super-resolution reconstruction. In the 0.78 mm human data, the ROVER-dMRI slice thickness (10 mm) is roughly 10*/*3.8 ≈ 2.6 times that of the matched gSlider-MB2 acquisition (3.8 mm) at nearly the same in-plane resolution, which accounts for the substantially higher SNR observed in Figure 3. Crucially, as shown in FOD results in Figure 5 and the DTI analysis in Figure S2, ROVER-dMRI is capable of further reducing the scan time by 65% (from 20-view to 13-view). The same principle holds for the 0.6 mm monkey data, where the ROVER-dMRI slice thickness is twice that of gSlider. Since SNR increases linearly with slice thickness but only as the square root of the number of signal averages, a two-fold thickness advantage corresponds to the SNR that gSlider would reach only with four averages, i.e. a four-fold increase in scan time. Thick-slice rotating-view acquisition therefore provides an efficient route to the high base SNR required for mesoscale diffusion imaging, which the multi-view INR then converts into isotropic high-resolution volumes. A second SNR gain comes from the reconstruction itself: because the shared multi-head INR encodes all diffusion directions with a single hash feature, it pools their common anatomy, so uncorrelated per-direction noise averages out while the individual heads preserve direction-specific contrast.

In the current work, we characterize the in-plane blurring using a 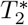-weighted phase-encode PSF simulation (Figure S3) ^38^, and note that this blurring is a property of the EPI readout itself, common to any EPI-based acquisition and not specific to, or introduced by, the ROVER-dMRI framework. More directly relevant to ROVER-dMRI is a slight blur that could likely arise from the reconstruction itself, akin to an intrinsic point-spread function. We quantified this trade-off directly using the 3D Laplacian variance (Section 2.4.2, Figure 4): where our ROVER-dMRI achieved a higher Laplacian variance than gSlider, indicating that its effective point-spread function remains sharper than that produced by the slab-deconvolution reconstruction of gSlider.

Despite these advancements, a limitation should be acknowledged: like other multi-view super-resolution strategies, ROVER-dMRI assumes consistent geometry across the rotated acquisitions, so inter-view subject motion can introduce misregistration. While we corrected for within view eddy current and motion artifacts, across view motion correction can be incorporated as demonstrated in Lyu et al. ^24^. Further, pretraining the representation, using priors learned across subjects or sessions rather than fitting each scan from scratch, could accelerate convergence and further stabilize reconstruction under motion and undersampling. Together with the open-source Pulseq sequence and reconstruction pipeline released with this work, these directions point toward practical mesoscale diffusion imaging in vivo.

## 5 CONCLUSION

We presented ROVER-dMRI, a unified rotating-view acquisition and super-resolution reconstruction framework for in vivo mesoscale diffusion MRI. Thick slices acquired across many rotated orientations significantly boost the base SNR, and a multi-resolution hash-encoded implicit neural representation jointly reconstructs the diffusion gradients into continuous isotropic volumes at up to 13× super-resolution without externally trained priors; a physics-informed forward model using Bloch-simulated slice-profile weights removed line artifacts and improved signal fidelity. Across human (0.78 mm and 0.5 mm) and monkey (0.6 mm) experiments on scanners with differing gradient performance, ROVER-dMRI delivered higher SNR than gSlider and the vendor product sequence, preserved coherent DEC and FOD orientations, and retained microstructural fidelity with the acquisition reduced from 20 to 13 views. Together with the open-source Pulseq sequence and reconstruction pipeline released with this work, ROVER-dMRI offers a practical route to high-SNR, whole-brain mesoscale diffusion imaging.

## ACKNOWLEDGMENTS

This study is supported by National Institutes of Health grants R01MH116173 and R01MH132610.

## SUPPORTING INFORMATION

**FIGURE S1.**
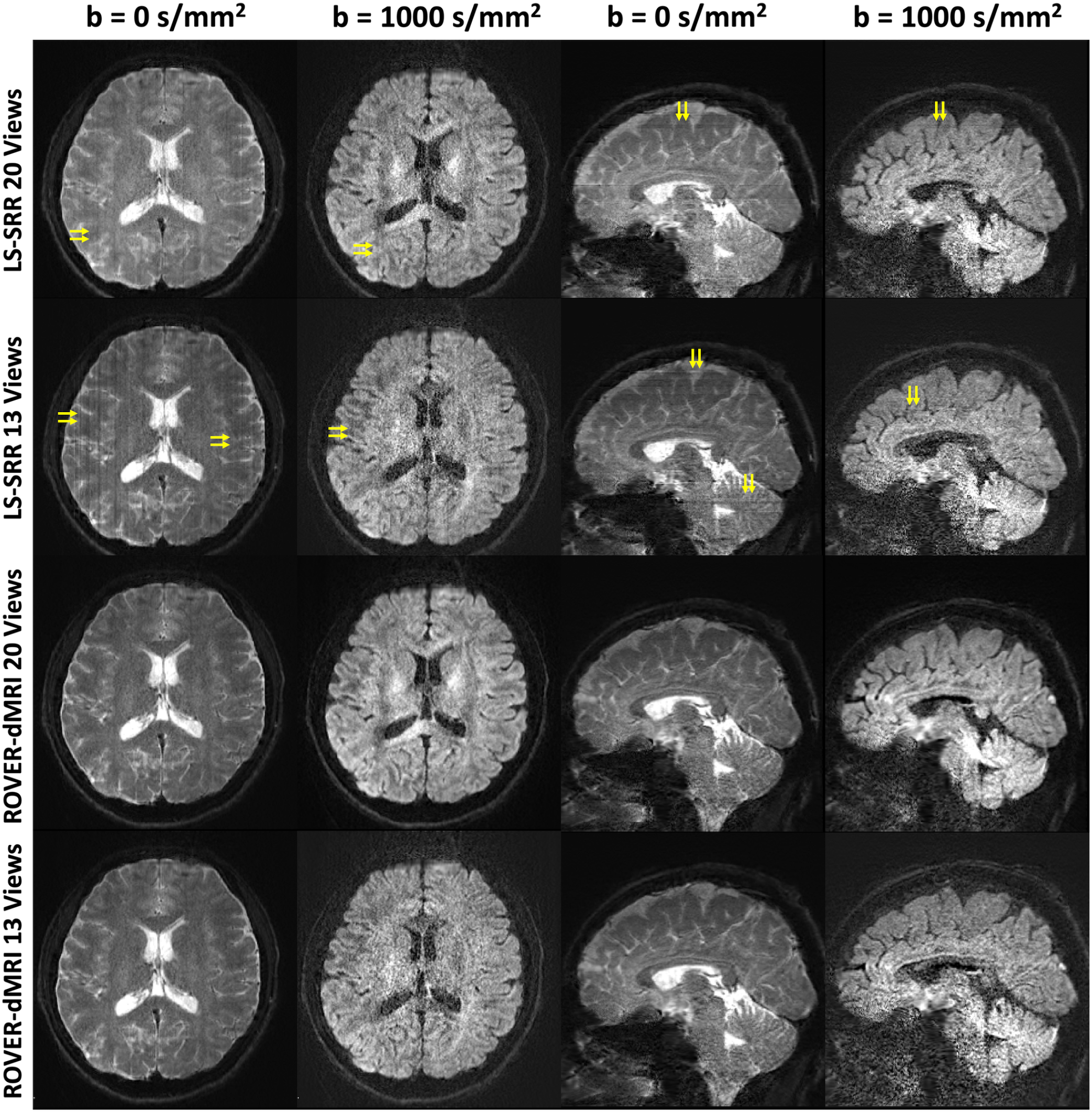
Representative axial and sagittal *b* = 0 and *b* = 1000 s*/*mm^2^ images comparing LS-SRR and ROVER-dMRI at 0.78 mm isotropic resolution. From top to bottom rows: LS-SRR with 20 views, LS-SRR with 13 views, ROVER-dMRI with 20 views, and ROVER-dMRI with 13 views. Yellow arrows indicate line artifacts in the LS-SRR reconstructions; these artifacts become more pronounced as the number of views is reduced from 20 to 13. ROVER-dMRI shows no such line artifacts.

**FIGURE S2.**
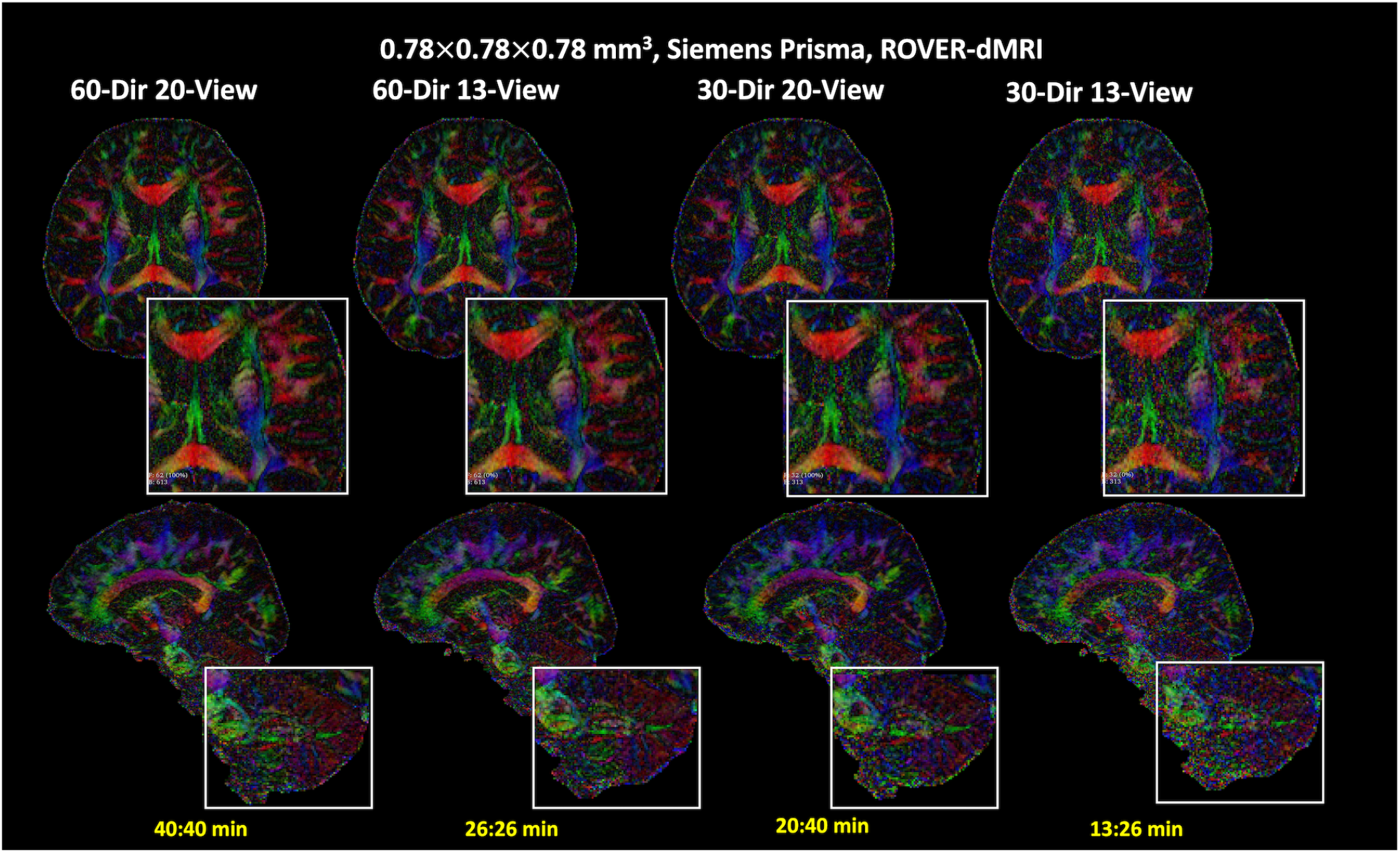
DEC fractional-anisotropy maps from ROVER-dMRI at 0.78 mm isotropic resolution (Siemens Prisma) for varying numbers of diffusion directions (60 or 30) and views (20 or 13), with the 60-direction, 20-view reconstruction serving as the reference. Top row: axial slice with a zoomed inset of the corona radiata. Bottom row: coronal slice with a zoomed inset of the cerebellum. White-matter structure in both regions remains clear when the number of diffusion directions is reduced from 60 to 30, while reducing the number of views from 20 to 13 introduces only a slight increase in noise. These results demonstrate that ROVER-dMRI is robust to the increased noise associated with a reduced number of diffusion directions, and that the joint reconstruction remains accurate even when the number of views (13) falls below the Nyquist sampling rate required by the reference 20-view acquisition.

**FIGURE S3.**
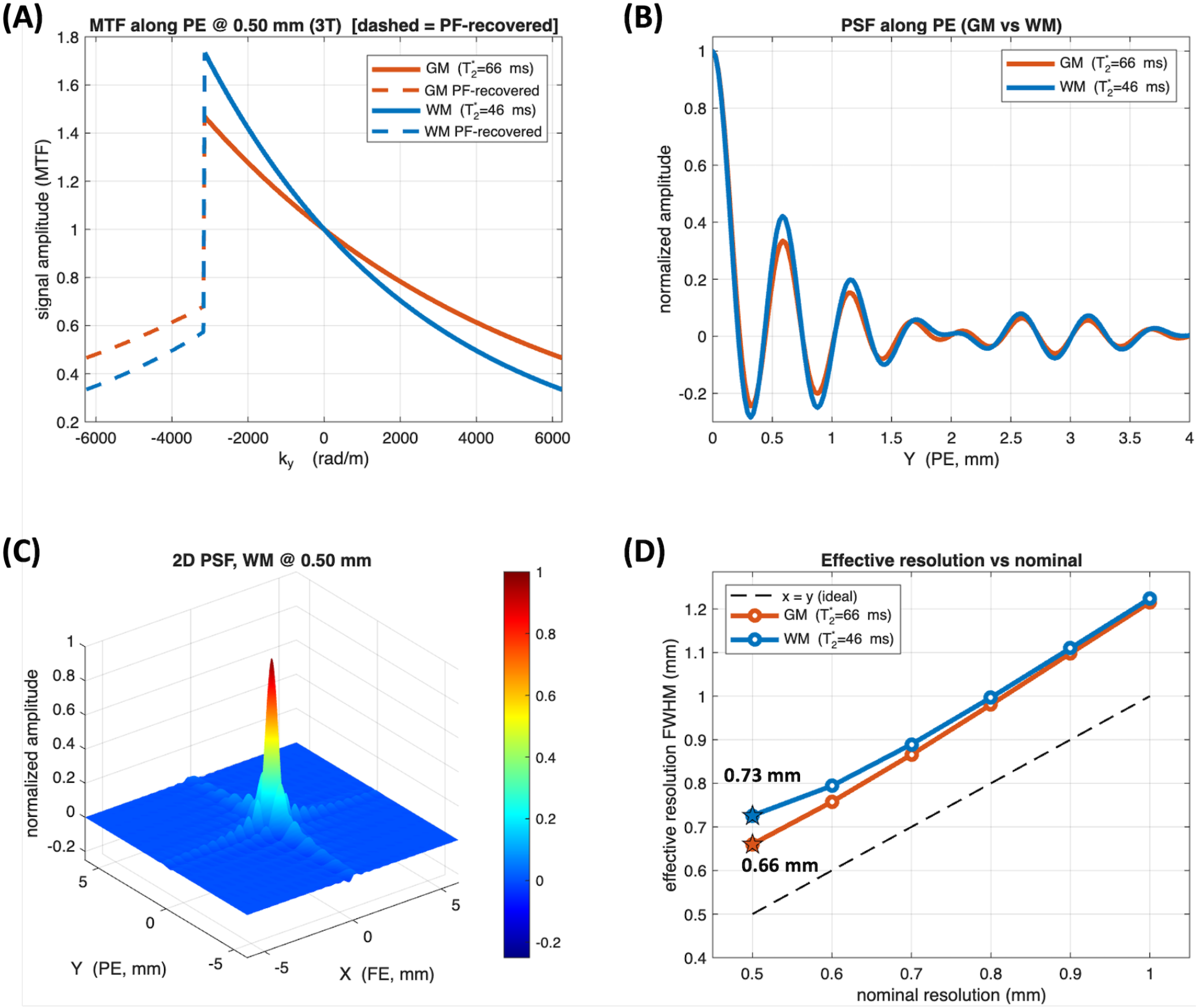
Point-spread-function analysis of the 0.5 mm 3-shot partial-Fourier EPI protocol (LORAKS reconstruction) at 3 T. (A) 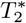 modulation transfer function (MTF) along the phase-encode (PE) direction for gray matter 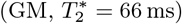 and white matter 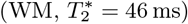, normalized to unity at *k*_*y*_ = 0. Solid lines are the acquired *k*-space data; dashed lines are the partial-Fourier region restored by conjugate (Hermitian) symmetry. The MTF falls off faster for WM because of its shorter 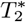. (B) Corresponding PE point-spread functions (real part) for GM and WM. The narrow main lobe is accompanied by side lobes (Gibbs ringing); WM shows larger side lobes and a broader profile. (C) Two-dimensional PSF for WM at the 0.5 mm nominal resolution. Blurring and ringing are concentrated along the PE axis, while the fully sampled readout (FE) axis is close to ideal. (D) Effective resolution (FWHM of the normalized-magnitude PE PSF) versus nominal resolution for GM and WM. The dashed line is the identity (*x* = *y*); stars mark the 0.5 mm operating point (GM 0.66 mm, WM 0.73 mm). Effective resolution exceeds the nominal pixel at all resolutions and diverges from the identity line as the nominal resolution becomes finer and the readout lengthens.

## REFERENCES

1. Miller Karla L., Alfaro-Almagro Fidel, Bangerter Neal K., et al. Multimodal population brain imaging in the UK Biobank prospective epidemiological study. Nature Neuroscience. 2016;19:1523–1536.

2. Johansen-Berg Heidi, Behrens Timothy E.J., Sillery Emma, et al. Functional-anatomical validation and individual variation of diffusion tractography-based segmentation of the human thalamus. Cerebral Cortex. 2005;15(1):31–39.

3. Misquitta Karen, Daou Marc, Conklin John, et al. Detecting silent acute microinfarcts in cerebral small vessel disease using submillimeter diffusion-weighted magnetic resonance imaging: Preliminary results. Stroke. 2022;53(7):e251–e252.

4. Song Allen W., Chang Hing-Chiu, Petty Christopher, et al. Improved delineation of short cortical association fibers and gray/white matter boundary using whole-brain three-dimensional diffusion tensor imaging at submillimeter spatial resolution. Brain Connectivity. 2014;4(9):636– 640.

5. Zhang Fan, Chen Yuqian, Ning Lipeng, et al. Assessment of the depiction of superficial white matter using ultrahigh-resolution diffusion MRI. Human Brain Mapping. 2024;45(14):e70041.

6. Reveley Colin, Seth Anil K., Pierpaoli Carlo, et al. Superficial white matter fiber systems impede detection of long-range cortical connections in diffusion MR tractography. Proceedings of the National Academy of Sciences. 2015;112(21):E2820–E2828.

7. Calabrese Evan, Hickey Patrick, Hulette Christine, et al. Postmortem diffusion MRI of the human brainstem and thalamus for deep brain stimulator electrode localization. Human Brain Mapping. 2015;36(8):3167–3178.

8. Ning Lipeng, Setsompop Kawin, Michailovich Oleg, et al. A joint compressed-sensing and super-resolution approach for very high-resolution diffusion imaging. NeuroImage. 2016;125:386–400.

9. Vis Geraline, Nilsson Markus, Westin Carl-Fredrik, et al. Accuracy and precision in super-resolution MRI: Enabling spherical tensor diffusion encoding at ultra-high b-values and high resolution. NeuroImage. 2021;245:118673.

10. Engström Mathias, Skare Stefan. Diffusion-weighted 3D multislab echo planar imaging for high signal-to-noise ratio efficiency and isotropic image resolution. Magnetic Resonance in Medicine. 2013;70(6):1507–1514.

11. Wu Wenchuan, Poser Benedikt A., Douaud Gwenäelle, et al. High-resolution diffusion MRI at 7T using a three-dimensional multi-slab acquisition. NeuroImage. 2016;143:1–14.

12. Dai Erpeng, Liu Simin, Guo Hua. High-resolution wholebrain diffusion MRI at 3T using simultaneous multi-slab (SMSlab) acquisition. NeuroImage. 2021;237:118099.

13. Li Ziyu, Zhu Silei, Miller Karla L., et al. Submillimeter diffusion MRI using an in-plane segmented 3D multislab acquisition and denoiser-regularized reconstruction. Medical Image Analysis. 2026;107:103834.

14. Setsompop Kawin, Fan Qiuyun, Stockmann Jason, et al. High-resolution in vivo diffusion imaging of the human brain with generalized slice dithered enhanced resolution: Simultaneous multislice (gSlider-SMS). Magnetic Resonance in Medicine. 2018;79(1):141–151.

15. Ramos-Llordén Gabriel, Ning Lipeng, Liao Congyu, et al. High-fidelity, accelerated whole-brain submillimeter in vivo diffusion MRI using gSlider-spherical ridgelets (gSlider-SR). Magnetic resonance in medicine. 2020;84(4):1781–1795.

16. Liao Congyu, Bilgic Berkin, Tian Qiyuan, et al. Distortion-free, high-isotropic-resolution diffusion MRI with gSlider BUDA-EPI and multicoil dynamic B0 shimming. Magnetic Resonance in Medicine. 2021;86(2):791– 803.

17. Jun Yohan, Liu Qiang, Gong Ting, et al. PRIME: Phase reversed interleaved multi-echo acquisition enables highly accelerated distortion-corrected diffusion MRI. Medical Image Analysis. 2026;111:104058.

18. Nicastro Michele, Jeurissen Ben, Beirinckx Quinten, et al. To shift or to rotate? Comparison of acquisition strategies for multi-slice super-resolution magnetic resonance imaging. Frontiers in Neuroscience. 2022;16:1044510.

19. Dong Zijing, Reese Timothy G., Lee Hong-Hsi, et al. Romer-EPTI: Rotating-view motion-robust superresolution EPTI for SNR-efficient distortion-free in-vivo mesoscale diffusion MRI and microstructure imaging. Magnetic Resonance in Medicine. 2025;93(4).

20. Sitzmann Vincent, Martel Julien N.P., Bergman Alexander W., et al. Implicit neural representations with periodic activation functions. Advances in Neural Information Processing Systems. 2020;33:7462–7473.

21. Xie Christopher, Park Keunhong, Martin-Brualla Ricardo, et al. Fig-NeRF: Figure-ground neural radiance fields for 3D object category modelling. In: Proceedings of the IEEE/CVF Conference on Computer Vision and Pattern Recognition (CVPR), :962–971; 2022.

22. Müller Thomas, Evans Alex, Schied Christoph, et al. Instant neural graphics primitives with a multiresolution hash encoding. ACM Transactions on Graphics (TOG). 2022;41(4):1–15.

23. Dwedari Mohammed Munzer, Consagra William, Müller Philip, et al. Estimating neural orientation distribution fields on high-resolution diffusion MRI scans. In: Medical Image Computing and Computer Assisted Intervention (MICCAI), :307–317; 2024.

24. Lyu Jun, Ning Lipeng, Consagra William, et al. Rapid whole brain motion-robust mesoscale in-vivo MR imaging using multi-scale implicit neural representation. Medical Image Analysis. 2026;107:103830.

25. Mugler John P.. Optimized three-dimensional fast-spinecho MRI. Journal of Magnetic Resonance Imaging. 2014;39(4):745–767.

26. Layton Kelvin J., Kroboth Stefan, Jia Feng, et al. Pulseq: A rapid and hardware-independent pulse sequence prototyping framework. Magnetic Resonance in Medicine. 2017;77(4):1544–1552.

27. Müller Thomas. tiny-cuda-nn https://github.com/NVlabs/tiny-cuda-nnVersion2.0, BSD-3-Clause license; 2021.

28. Rosu Radu Alexandru, Behnke Sven. PermutoSDF: Fast Multi-View Reconstruction with Implicit Surfaces Using Permutohedral Lattices. In: Proceedings of the IEEE/CVF Conference on Computer Vision and Pattern Recognition (CVPR), :8466–8475; 2023.

29. Plenge Esben, Poot Dirk H.J., Bernsen Monique, et al. Super-resolution methods in MRI: Can they improve the trade-off between resolution, signal-to-noise ratio, and acquisition time?. Magnetic Resonance in Medicine. 2012;68(6):1983–1993.

30. Liu Qiang, Ning Lipeng, Shaik Imam Ahmed, et al. Reduced cross-scanner variability using vendor-agnostic sequences for single-shell diffusion MRI. Magnetic Resonance in Medicine. 2024;92(1):246–256.

31. Liu Qiang, Gagoski Borjan, Shaik Imam Ahmed, et al. Time-division multiplexing (TDM) sequence removes bias in T2 estimation and relaxation-diffusion measurements. Magnetic Resonance in Medicine. 2024;92(6):2506–2519.

32. Andersson Jesper L.R., Sotiropoulos Stamatios N.. An integrated approach to correction for off-resonance effects and subject movement in diffusion MR imaging. NeuroImage. 2016;125:1063–1078.

33. Inati Souheil J., Naegele Joseph D., Zwart Nicholas R., et al. ISMRM Raw data format: A proposed standard for MRI raw datasets. Magnetic Resonance in Medicine. 2017;77(1):411–421.

34. Hoge W. Scott, Polimeni Jonathan R.. Dual-polarity GRAPPA for simultaneous reconstruction and ghost correction of echo planar imaging data. Magnetic Resonance in Medicine. 2016;76(1):32–44.

35. Jiang Yuancheng, Jun Yohan, Liu Qiang, et al. Multishot dual-polarity GRAPPA: Robust Nyquist ghost correction for multishot EPI. Magnetic Resonance in Medicine. 2026;95(5):2726–2736.

36. Wang Nan, Abraham Daniel, Shah Zahra, et al. Field-correcting GRAPPA (FCG): A technique to correct spatiotemporal-varying phase errors in echo planar imaging. bioRxiv. 2026;.

37. Haldar Justin P.. Low-rank modeling of local k-space neighborhoods (LORAKS) for constrained MRI. IEEE Transactions on Medical Imaging. 2014;33(3):668–681.

38. Feizollah Sajjad, Tardif Christine L.. High-resolution diffusion-weighted imaging at 7 Tesla: Single-shot readout trajectories and their impact on signal-to-noise ratio, spatial resolution and accuracy. NeuroImage. 2023;274:120159.

39. Peters Andrew M., Brookes Matthew J., Hoogenraad Frank G., et al. T2* measurements in human brain at 1.5, 3 and 7 T. Magnetic Resonance Imaging. 2007;25(6):748– 753.

40. Cox Emma F., Gowland Penny A.. Simultaneous quantification of T2 and T2′ using a combined gradient echo-spin echo sequence at ultrahigh field. Magnetic Resonance in Medicine. 2010;64(5):1441–1450.

41. Pech-Pacheco José Luis Cristóbal Gabriel, Chamorro-Martínez Jesús, et al. Diatom autofocusing in brightfield microscopy: a comparative study. In: Proceedings 15th International Conference on Pattern Recognition (ICPR-2000), :314–317; 2000.

42. Wang Fuyixue, Dong Zijing, Tian Qiyuan, et al. In vivo human whole-brain Connectom diffusion MRI dataset at 760 µm isotropic resolution. Scientific Data. 2021;8:122.

