## Supplementary figures and images for "High-SNR Whole-Brain Mesoscale Diffusion MRI Using Rotating-View Acquisitions (ROVER-dMRI)"

### Supporting Information Figure 1

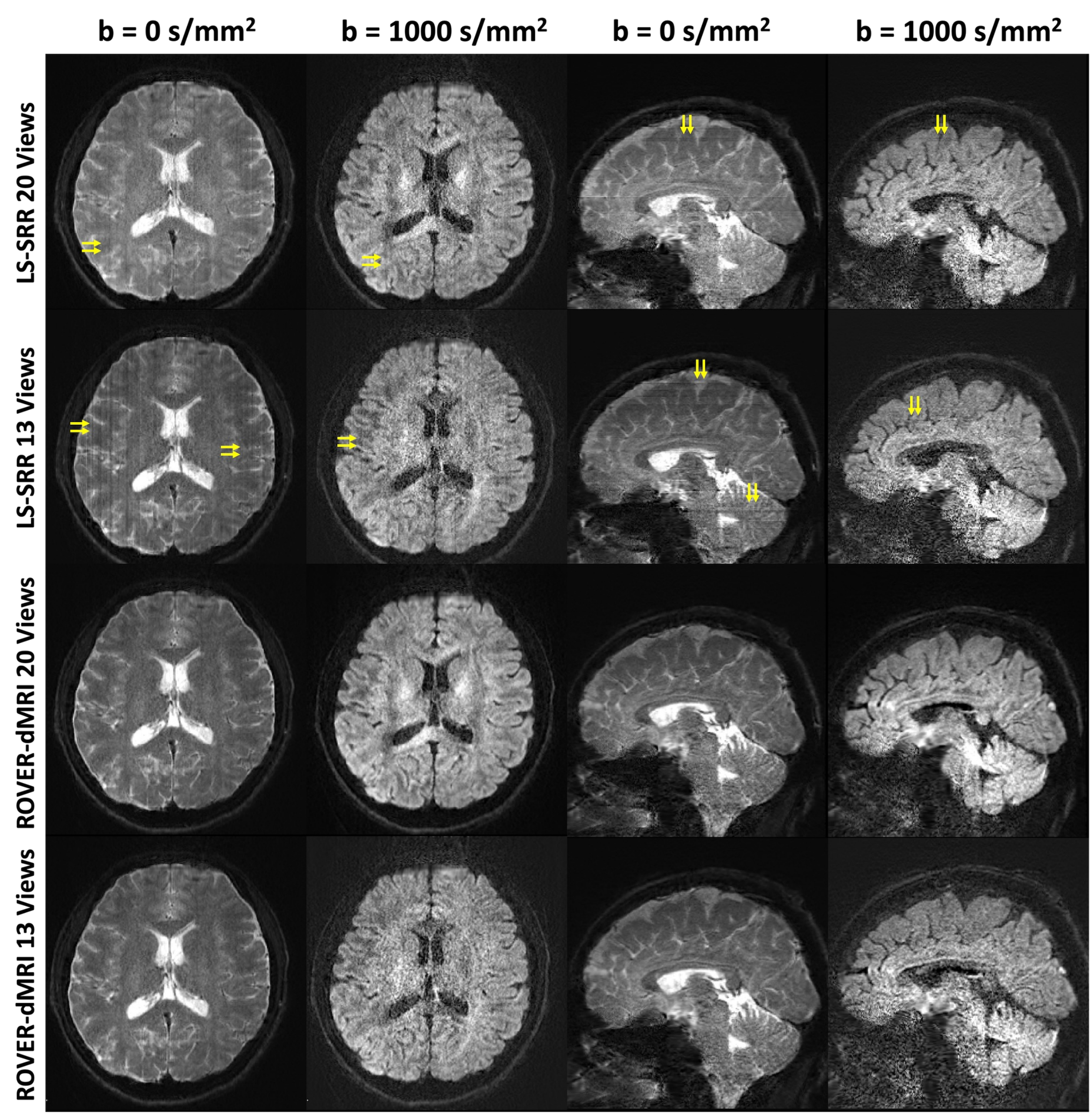

### Supporting Information Figure 2

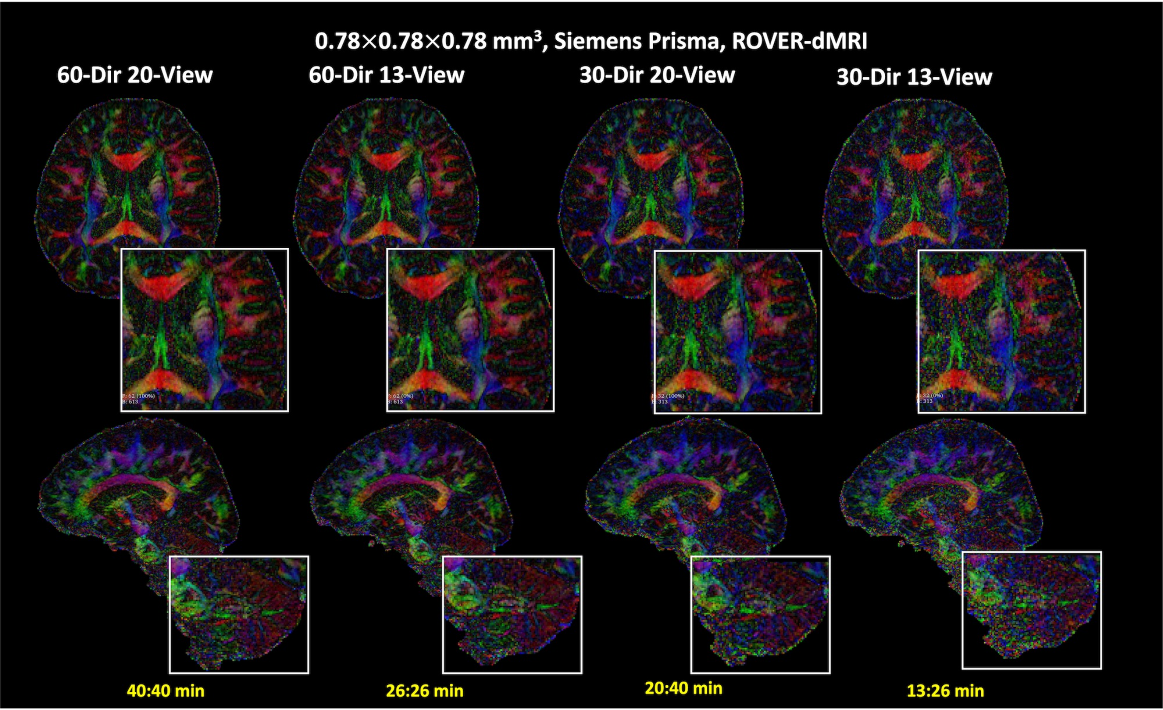

### Supporting Information Figure 3

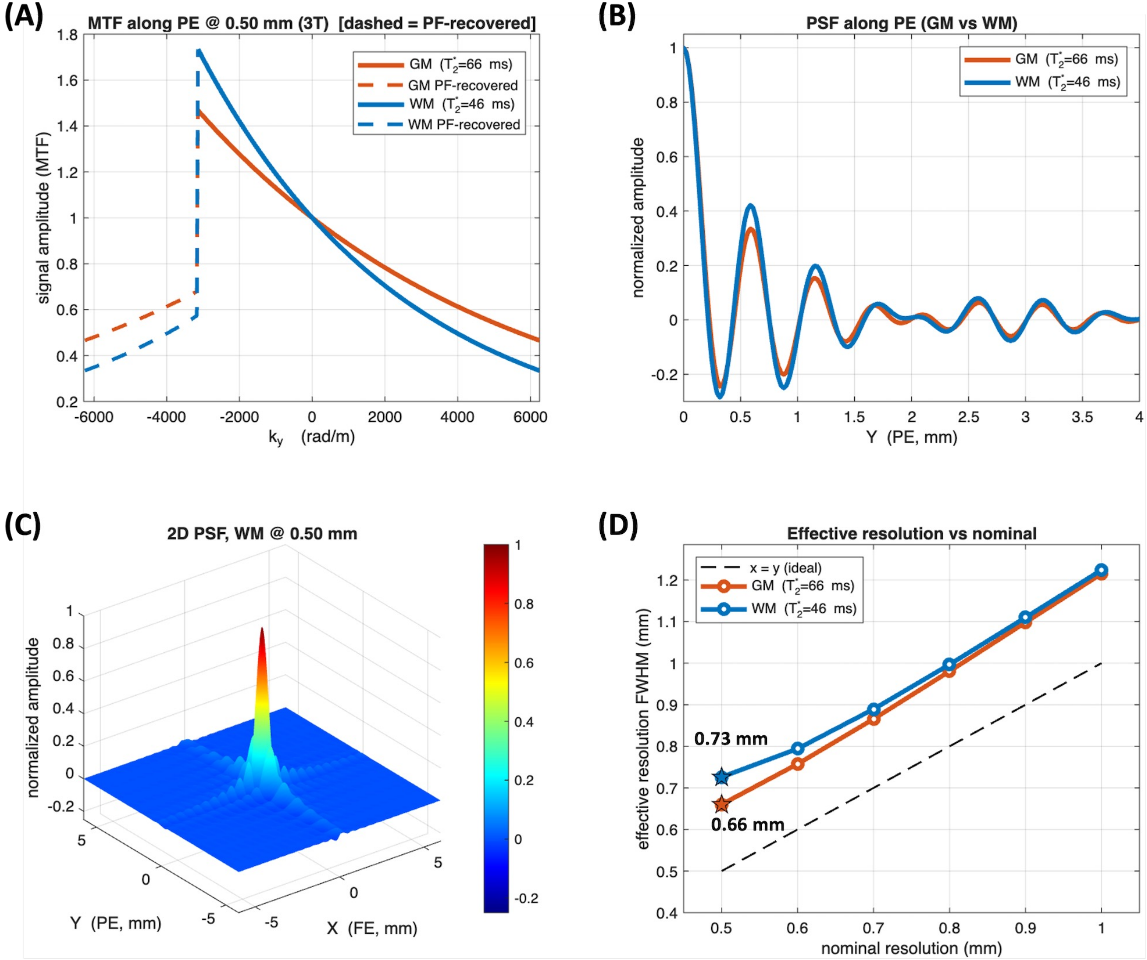
